# Cardiomyocyte autophagy promotes a pro-regenerative immune response during cardiac regeneration

**DOI:** 10.64898/2026.03.16.712191

**Authors:** Florian Constanty, Bailin Wu, Srishti Shekhar, Alina Bektimirova, Vasiliki Bakali, Lucia Blasco Almodovar, Frauke Senger, Norbert Frey, Arica Beisaw

**Author notes:** These authors contributed equally (Florian Constanty, Bailin Wu).

## Abstract

Adult zebrafish possess a remarkable ability to regenerate their heart following cardiac injury. Over the past decades, our understanding of the diverse cell types involved in zebrafish cardiac regeneration has greatly advanced. However, the mechanisms governing their interaction and how heterocellular crosstalk drives regeneration remain poorly understood. Here, we identify cardiomyocyte autophagy as a key link between the cardiomyocyte injury response and heterocellular crosstalk between cardiomyocytes and macrophages. We find that cardiomyocyte autophagy is downstream of AP-1 transcription factors. Using newly generated genetic tools, we find that cardiomyocyte autophagy is an important regulator of cardiomyocyte protrusion into the fibrotic injured tissue and its disruption leads to defects in scar resolution. Notably, we find that blocking cardiomyocyte autophagy has a marked effect on the transcriptomic signatures in cardiac macrophages, shifting gene expression from phagocytic/pro-inflammatory/pro-reparative towards pro-angiogenic and pro-fibrotic states. Altogether, our results uncover autophagy as a mechanism linking cardiomyocyte injury responses to macrophage phenotype and coordinated tissue remodeling during heart regeneration.

## INTRODUCTION

Despite substantial advances in the field of cardiovascular biology, heart disease globally remains the leading cause of illness and death^1,2^. Following an acute myocardial infarction in the human heart, millions of cardiomyocytes (CMs) are lost and replaced with a fibrotic scar. While this scar is necessary to preserve the structural integrity of the heart, its presence, combined with reduced contractile CM mass, ultimately contributes to the progression towards heart failure. There remains a critical need to develop and refine therapeutic strategies capable of restoring lost cardiac tissue. In contrast to mammals, non-mammalian vertebrates such as teleost zebrafish can regenerate their hearts following various types of injury, including resection^3^, genetic ablation of CMs^4^, and cryoinjury^5–7^. Cryoinjury of the apex of the heart leads to cell death, activation of an inflammatory response, and deposition of a fibrotic collagen-containing scar, similar to myocardial infarction in humans. Although research in the past two decades has substantially expanded our understanding of the multiple cell types involved in cardiac regeneration in zebrafish, less is known about the mechanisms underlying their interaction and how heterocellular crosstalk promotes cardiac regeneration.

We have previously shown that AP-1 transcription factors (TFs) are essential regulators of the CM regeneration process through their ability to promote changes in the chromatin accessibility landscape^8^. Blocking CM AP-1 function with dominant-negative A-Fos led to defects in CM dedifferentiation, proliferation, and protrusion into the injured tissue, and ultimately resulted in the remaining presence of a fibrotic scar at 90 days post cryoinjury^8^. Notably, utilizing similar methods with dominant negative A-Fos in epicardial-derived and endocardial cells, others have shown that AP-1 plays an essential role downstream of Tnfa to promote the expression of fibroblast-specific genes, presence of epicardial-derived cells at the wound, and endocardial activation^9^. The effects in fibroblasts were shown to be mediated by AP-1’s function in regulating H3K27ac at putative enhancer regions of genes, including *fn1a, acta2,* and *col5a3b*.

While AP-1 plays a clear role in regulating chromatin accessibility and gene expression programs in multiple cell types during cardiac regeneration, AP-1 TFs are also well-known for their role in regulating the response to cellular stress. Systems biology approaches revealed AP-1 transcription factors as potential upstream regulators of genes involved in autophagy/lysosomal function and regulation^10^. However, further functional studies have revealed that regulation of autophagy by AP-1 is highly context-dependent. For example, while Jun and JunB have been shown to inhibit starvation-induced autophagy in HeLa cells^11^, Jun has been shown to promote autophagy induced by ceramide in cancer cells through transcriptional upregulation of genes encoding positive regulators of autophagy, including LC3 and Beclin-I^12,13^. In the heart, Fosl2 has been shown to promote autophagy in cardiac fibroblasts, resulting in cardiac fibrosis and pathological remodeling of the heart in response to cardiac hypertrophy^14^.

Autophagy is a highly regulated process used by the cell to degrade or recycle damaged proteins, organelles, or cytoplasmic components in response to stress and relies on a set of proteins known as autophagy-related (Atg) proteins^15^. Autophagy is initiated under conditions of cellular stress and involves the activation of the Atg1/ULK complex, which triggers formation of the isolation membrane. Membrane nucleation and expansion are coordinated by several Atg proteins, including the Atg12-Atg5-Atg16 complex, which functions in a ubiquitin-like conjugation system essential for autophagosome formation. A key component of the pathway is LC3 (the vertebrate homolog of yeast Atg8), which is proteolytically cleaved by Atg4 in the cytosol (LC3-I) and becomes lipidated with phosphatidylethanolamine to form LC3-II. LC3-II incorporates into the growing phagophore membrane and is important for cargo selection, and phagophore membrane expansion/closure^16,17^. Phagophore membrane closure and fusion of the autophagosome to the lysosome results in degradation and recycling of damaged proteins and organelles for reuse by the cell.

The role of autophagy in CMs is largely considered cardioprotective^18^, but many groups have shown that this role is likely dependent on context: for example, CM autophagy has been shown to play divergent roles depending on cardiac disease/injury model and the level and duration of autophagy^19–22^. In cardiac regeneration in zebrafish, there is a clear lack of consensus on the role of autophagy following cardiac injury. While two studies report an increased presence of autophagosomes in the heart following cardiac cryoinjury, they differ in their conclusions regarding whether autophagy accelerates or inhibits cardiac regeneration^23,24^. Furthermore, the role of autophagy specifically in CMs in cardiac regeneration in zebrafish remains unexplored. Here, we show that AP-1 promotes cardiomyocyte autophagy in response to cardiac cryoinjury in zebrafish and developed genetic tools to block autophagy specifically in CMs. Blocking CM autophagy did not have dramatic effects on cell survival, dedifferentiation or proliferation in response to injury. However, CM protrusion and replacement of the fibrotic scar were affected by the lack of CM autophagy. Strikingly, blocking CM autophagy had a significant effect on the phenotype of cardiac macrophages, skewing the macrophage transcriptome from a pro-inflammatory/pro-reparative gene expression signature to a pro-fibrotic/angiogenic signature, suggesting that CM autophagy can mediate heterocellular crosstalk between CMs and macrophage in the zebrafish heart.

## METHODS

### Animal Experimentation

All animal experimentation in this study was performed in compliance with EU Directive 2010/63/EU and national ethical and animal welfare guidelines (RP Karlsruhe). Zebrafish lines used in this study include: *Tg(mpeg1:EGFP)gl22*^25^, *Tg(cryaa:DsRed, −5.1myl7:CreERT2)pd10*^26^, *Tg(p14a.ubbR:loxp-STOP-loxp-FlagAFos-P2A-TagBFPHA, gcry:mBFP)* (this study) and *Tg(p14a.ubbR:loxp-STOP-loxp-3xHAAtg4bC74A, gcry:mBFP)* (this study), all of which harbored a wild-type AB genetic background.

### Generation of transgenic lines

Plasmids constructed in this study were cloned using conventional restriction enzymes (New England Biolabs), NEB 5-alpha Competent E. coli High Efficiency (New England Biolabs, C2987) and NEBuilder HiFi DNA Assembly Master Mix (New England Biolabs, E2621). To construct the A-Fos vector for insertion into the pIGLET14a locus, the following fragments were PCR amplified and assembled via Gibson assembly: *ubbR* promoter^27^, loxP-STOP-loxP cassette^28^, Flag-Afos-P2A-TagBFP-HA-SV40 cassette^8^, with a 3’-5’ Ocean Pout Terminator and 3’-5’ gcry promoter-NLS-monomeric BFP cassette^29^ (a gift from Jeffrey Essner, Addgene plasmid #117765). An attB site (GATGGGTGAGGTGGAGTACGCGCCCGGGGAGCCCAAGGGCACGCCCTGGCACCCGCACCGCGGCTTCGAG) was cloned upstream of the *ubbR* promoter and bacterial vector sequence was flanked by minimal FRT sites. To construct the Atg4bC74A vector for insertion into the pIGLET14a locus, 3xHA-Atg4bC74A was inserted into the abovementioned pIGLET-AFos backbone by replacing the Flag-Afos-P2A-TagBFP cassette. The first fragment (encoding amino acids 1-73) of *atg4b* was amplified by PCR from zebrafish 3-day post fertilization (dpf) embryonic cDNA, with the 3xHA tag introduced via tailed PCR. The second fragment (encoding amino acids 74-394) was also amplified from zebrafish 3-dpf embryonic cDNA and contained the cysteine-to-alanine mutation at amino acid 74 (GCA) introduced via tailed PCR.

25 pg plasmid construct and 15 pg *phiC31* integrase mRNA were co-injected into one-cell stage pIGLET14a embryos^30^ to generate *Tg(p14a.ubbR:loxp-STOP-loxp-FlagAFos-P2A-TagBFP-HA, gcry:mBFP) and Tg(p14a.ubbR:loxp-STOP-loxp-3xHAAtg4bC74A, gcry:mBFP)* lines. The pT3TS-phiC31 integrase plasmid (a gift from Thomas Juan and Didier Stainier) was linearized with KpnI and transcribed *in vitro* using the T3 mMessage mMachine Kit (Thermo Fisher Scientific, AM1348) according to the manufacturer’s instructions. F1 animals were genotyped with the following primers to confirm targeted integration: forward, 5’-CATGATCGAAAACGAGCAATGT-3’; reverse, 5‘-CGATTAAGTTGGGTAACGCCAGG-3’.

### Cryoinjury of zebrafish hearts and induction of transgene expression

Cryoinjury was performed as previously described^5–7^. Adult zebrafish (4–12 months old) were anesthetized in 0.025% Tricaine, and a small incision was made through the skin and pericardial sac to expose the heart. A metal probe cooled in liquid nitrogen was then applied to the ventricular apex. Following the procedure, fish were placed in system water to recover.

For the induction of CM-specific overexpression of *A-Fos* and *atg4bC74A*, adult *Tg(p14a.ubbR:loxp-STOP-loxP-FlagAFos-P2A-TagBFPHA); Tg(myl7:CreERT2)* (referred to as CM:AFos) and *Tg(p14a.ubbR:loxp-STOP-loxP-3xHAAtg4bC74A); Tg(myl7:CreERT2)* (referred to as CM:Atg4bC74A) zebrafish were anesthetized in 0.025% Tricaine and 10 μL of 0.5 mg/mL of 4-hydroxytamoxifen (4-HT) was injected intraperitoneally for 3 consecutive days before cryoinjury. As a control, Cre(-) *Tg(p14a.ubbR:loxp-STOP-loxP-FlagAFos-P2A-TagBFPHA)* and *Tg(p14a.ubbR:loxp-STOP-loxP-3xHAAtg4bC74A)* were anesthetized in 0.025% Tricaine and 10 μL of ethanol vehicle was injected intraperitoneally for 3 consecutive days before cryoinjury. For immunoblotting experiments, adult CM:AFos and CM:Atg4bC74A zebrafish were anesthetized in 0.025% Tricaine and 10 μL of 0.5 mg/mL of 4-hydroxytamoxifen (4-HT) or ethanol vehicle was injected intraperitoneally for 3 consecutive days. As a further control, Cre(-) *Tg(p14a.ubbR:loxp-STOP-loxP-FlagAFos-P2A-TagBFPHA)* and *Tg(p14a.ubbR:loxp-STOP-loxP-3xHAAtg4bC74A)* were anesthetized in 0.025% Tricaine and 10 μL of 0.5 mg/mL of 4-hydroxytamoxifen (4-HT) was injected intraperitoneally for 3 consecutive days.

### Immunostaining, imaging, and quantification

Adult zebrafish hearts were harvested and treated as previously described^31^. For immunostaining of Mef2/PCNA, samples were treated as previously described^32^. For Lc3b and Sqstm1(p62) immunostaining, sections were permeabilized by incubation in acetone for 15 min at −20 °C. For immunostaining of Mpx and embCMHC (N2.261), and staining using the CNA35 probe, antigen retrieval with sodium citrate (pH 6.0) at 95°C for 20 min was required. Primary antibodies used in this study include: anti-MHC-Clone A4.1025 (Merck 05-716-I-100UL) at 1:200, anti-Lc3b (Cell Signaling 2775) at 1:200, anti-SQSTM1/P62 (Genetex GTX100685) at 1:200, anti-GFP (Aves Labs GFP-1020) at 1:500, anti-Mpx (Genetex GTX128379) at 1:200, anti-embCMHC-Clone N2.261 (Santa Cruz sc-53096) at 1:100, anti-Mef2 (Boster DZ01398-1) at 1:200, anti-PCNA (Abcam ab29) at 1:200, anti-Tnfa (Abcam ab1793) at 1:50, and anti-Cxcr4b (Abcam ab229623) at 1:250. Alexa Fluor-coupled secondary antibodies were used (Thermo Fisher) at 1:500. Other dyes and probes used in this study include CNA35 at 1:100 and Alexa Fluor 647-Phalloidin (Thermo Fisher A22287) at 1:300.

Imaging of immunostained sections was performed using a Leica Mica confocal microscope with 20× HC PL FLUOTAR and 63× HC PL APO objectives. Quantification of Lc3b and p62 was performed from images taken with a 63x objective at border zone or remote zone CMs. Lc3b signal was thresholded and particles with a size more than 5 pixels were counted per frame. p62 signal was quantified by mean intensity per frame. Quantification of *mpeg1*:EGFP+ cells was performed from cryosections in the area 50 μm distal and 50 μm proximal to the wound border zone in three non-consecutive sections exhibiting the largest wound area per heart. Quantification of mean intensity of embCMHC (N2.261) was performed in the CM area 100 μm proximal to the wound border zone. Particle counting of Mef2 and PCNA was performed in the CM area 100 μm proximal from the wound border in three nonconsecutive sections exhibiting the largest injured area from each heart. Quantification of mean intensity of CNA35 was performed from cryosections in the area 50 μm distal and 50 μm proximal to the wound border zone. Quantification of Cxcr4b mean intensity was performed within the entire injured area and in the CM area 100 μm proximal to the border zone. Quantification of mean intensity of Tnfa and Mpx was performed within the wound area. Mpx signal was thresholded and particles with a size more than 16 pixels were counted in the entire injured area and in the CM area 100 μm proximal to border zone and normalized to the quantified area. For measurements and quantification of cardiomyocyte protrusion at the border zone, 60 μm thick cryosections were obtained, and quantification was performed as previously described^31^. All quantifications described above were performed in ImageJ (v2.16.0).

### TUNEL assay, imaging, and quantification

For *in situ* cell death detection, sample preparation and TUNEL labeling were performed following manufacturer’s instructions (Roche, 11684795910). Imaging was performed with Leica Mica using confocal imaging, with a 20× HC PL FLUOTAR objective. TUNEL and DAPI signals were thresholded and particles with a size above 16 pixels were counted in the wound area in ImageJ (v2.16.0).

### Histological staining, imaging, and quantification

For Picrosirius Red staining, cryosections were incubated in Bouin’s solution 2 hrs at 56°C and then 1hr at room temperature, followed by a rinse in running water for 10 min. Sections were incubated in Picrosirius Red solution for 90 min at room temperature and subsequent steps were performed according to manufacturer’s instructions (Morphisto). Stained sections were mounted in Entellan (Sigma). Imaging was performed with Leica Mica using widefield imaging, with a 20× HC PL FLUOTAR objective. Scar area was quantified from three non-consecutive sections per heart relative to the ventricle area in ImageJ (v2.16.0).

### Hybridization chain reaction staining, imaging and quantification

mRNA expression of *mrc1b* and *col1a1a* was verified by HCR^TM^ RNA fluorescence *in situ* hybridization (RNA-FISH) as previously described^31^. Immunostaining was then performed according to the protocol above. Quantification of *mrc1b* mean intensity was performed from cryosections in the entire injured area and in the CM area 100 μm proximal to border zone in three nonconsecutive sections exhibiting largest wound per heart. For quantification of *col1a1a*, *mpeg1*:EGFP signal was thresholded as a mask and the mean intensity of GFP+ *col1a1a*+ signal was quantified in the entire injured area and in the CM area 100 μm proximal to border zone in two to three nonconsecutive sections exhibiting the largest wound per heart. A ratio of *col1a1a*+ GFP+ area to total GFP+ area was calculated in the entire injured area and in the CM area 100 μm proximal to border zone in two to three nonconsecutive sections exhibiting largest wound per heart. All quantifications were performed in ImageJ (v2.16.0).

### Culture of neonatal rat ventricular myocytes

Neonatal rat ventricular myocytes (NRVMs) were isolated from postnatal day 1-3 Sprague Dawley rat pups using the Neonatal Heart Dissociation kit, mouse/rat (130-098-373, Miltenyi) and the Neonatal Cardiomyocyte Isolation kit, rat (130-105-420, Miltenyi) according to manufacturer’s instructions, with the following adaptations: Briefly, collected hearts were washed in 1x ADS buffer to remove excess blood and placed in a petri dish on ice. Remaining blood vessels and non-cardiac tissues were carefully removed. Hearts were minced into small pieces and evenly distributed into C-tubes (approximately 20 hearts per tube). Enzyme Mix I and Enzyme Mix II were combined according to the manufacturer’s instructions, and 5 mL of the enzyme mixture was added to each C-tube. Tubes were tightly capped and placed on the GentleMACS dissociator for enzymatic digestion at 37 °C for 56 minutes. Following digestion, the tubes were briefly centrifuged at 1000 rpm at room temperature. To stop the enzymatic reaction, 7.5 mL of DMEM complete medium (P04_03600, PanBiotech) with 10% FCS (P04_03600, PanBiotech or FBS-11A, Capricorn Scientific), and 1% penicillin streptomycin (15140-122, Gibco) was added to each C-tube. The resulting cell suspension was passed through a 70 μm cell strainer placed on a 50 mL falcon tube pre-wetted with 2 mL DMEM complete medium. For NRVM isolation, the supernatant was carefully discarded, and the pellet was resuspended in 60 μl PEB buffer, 20 μl of isolation cocktail and 20 μl anti-red blood cell microbeads as per the manufacturer’s protocol. As a modification of the protocol, the cell suspension volume was adjusted to 20 mL later and centrifuged at 1000 rpm for 5 minutes at room temperature. The pellet was washed once more with 20 mL DMEM complete medium and centrifuged again under the same conditions. After the final wash, the supernatant was discarded and the NRVMs were resuspended in 30 mL DMEM complete medium. Cell number and viability were determined using a Countess automated cell counter by mixing the cell suspension with trypan blue at a 1:1 ratio. Finally, the cells were plated and incubated at 37 °C in a humidified incubator with 5% CO₂.

To test whether pharmacological mimicking of hypoxia can stimulate autophagy, NRVMs were treated with cobalt chloride (CoCl_2_, 8025400010, Sigma Aldrich) at a final concentration of 250 μM and chloroquine (CQ, C6628, Sigma Aldrich) at a final concentration of 10 mM, or a combination of CoCl_2_ and chloroquine at a final concentration of 250 μM and 10 mM, respectively, for 4 hours. To pharmacologically block AP-1 function, NRVMs were cultured in DMSO control or T-5224 (HY-12270, MedChemExpress) at a final concentration of 10 μM for a total of 24 hours. After 20 hours of pretreatment, CQ, CoCl_2_, and CQ+CoCl_2_ were added to the culture at the concentrations described above for a total of 4 hours.

### Immunoblotting

Whole cell lysates or whole adult zebrafish ventricles were collected and prepared for immunoblotting in 1X RIPA buffer (ab156034, Abcam) supplemented with cOmplete EDTA-free protease inhibitor cocktail (11873580001, Roche). Immunoblotting was performed per the standard protocol provided by BioRad, including the following adaptations. 35 μg of protein samples were loaded on SDS-PAGE and separated at 80V. The samples were then transferred to a 0.45 μm PVDF membrane at 550 mA for 90 minutes or 600 mA for 120 minutes for a low molecular weight protein or a high molecular weight protein of interest, respectively. After transfer the membrane was blocked with 5% skimmed milk for 1 h at room temperature. The blots were washed 3 times with 1X TBST for 3 minutes. After washing, the blots were incubated with primary antibodies for 12-14 hours at 4° C. Next, the blots were later washed 3 times with 1X TBST for 3 minutes, followed by probing with the respective HRP conjugated secondary antibodies for 1 h at room temperature. Subsequently, the blots were washed 3 times with 1X TBST for 3 minutes after probing with secondary antibodies. Finally, the blots were developed using Luminol or Femto (Thermo Scientific) and the bands were quantified using ImageJ (v2.16.0). Primary antibodies used in this study include: anti-HIF-1α (sc-13515, Santa Cruz) at 1:500, anti-JUN (166540, Santa Cruz) at 1:500, anti-FOSL2 (G-5, Santa Cruz) at 1:500, anti-LC3A (NB100-2331, Novus Biologicals) at 1:500, anti-Vinculin (VCL, V9131, Sigma Aldrich) at 1:1000, anti-beta-Actin (ACTB, sc-47778, Santa Cruz) at 1:500, anti-Flag (F1840, Sigma Aldrich) at 1:500, anti-HA (ab9110, Abcam) at 1:500. Secondary antibodies used in this study include: anti-rabbit IgG-HRP (7074P2, Cell Signaling Technology) at 1:5000, anti-mouse IgG-HRP (7076P2, Cell Signaling Technology) at 1:5000.

### Cardiomyocyte isolation from zebrafish ventricles

6 ventricles from control or CM:AFos fish at 7 dpci were isolated per biological replicate. Tissue dissociation using the Neonatal Heart Dissociation kit (Miltenyi 130-098-373) and cardiomyocyte isolation by sucrose density gradient centrifugation were performed as previously described^8,31^.

### Macrophage isolation from zebrafish ventricles

To isolate macrophages, three ventricles from control or CM:Atg4b fish at 7 dpci were used per biological replicate. Tissue dissociation was performed as previously described^32^. Isolated cells were filtered through a 30 μm nylon mesh and placed in ice-cold fresh FACS buffer (10% FBS and 1 mM EDTA in PBS). All subsequent steps were performed on ice and at 4 °C using DNA low-binding tubes (30108418, Eppendorf). Cells were washed twice with FACS buffer at 200 × g for 5 min and incubated in antibody blocking buffer (1× PBS, 1% DMSO, 2% normal goat serum) for 45 min in the dark with rotation. The macrophage population was identified by immunostaining for 30 min in the dark with rotation using an antibody against Csfr1 (rabbit, GTX128677, 0.75 μg/100 μl). Cells were then washed twice with FACS buffer and incubated with the secondary antibody (Alexa Fluor–conjugated α-rabbit IgG (H+L), raised in goat, Invitrogen, A11034) at 0.25 μg/100 μl for at least 20 min in the dark with rotation. Following incubation, cells were washed twice with FACS buffer at 200 × g for 5 min, and the pellet was resuspended in FACS buffer. Flow cytometry detection was started immediately. GFP⁺ cells were sorted using a Sony SH800S Cell Sorter equipped with a 100 μm nozzle. GFP fluorescence was measured using 30 mW 488 nm excitation with a 525/50 nm band-pass filter. Example plots showing the hierarchical gating strategy are provided in Supplementary Figure 5A. Macrophages were sorted directly into RLT buffer, stored at −80 °C until further processing, and subsequently used for RNA extraction.

### RNA extraction and RNA-sequencing library preparation

Isolated zebrafish cardiomyocytes were lysed in TRIzol (Invitrogen, 15596018). RNA was extracted using the TRIzol chloroform protocol according to the manufacturer’s instructions, followed by isopropanol precipitation in the presence of GlycoBlue (ThermoFisher, AM9515). RNA was resuspended in water and further purified using the RNA Clean & Concentrator kit (Zymo, R1019). RNA quality was assessed using TapeStation (Agilent). RNA from FACS-sorted macrophages was extracted using the Qiagen RNeasy Micro Kit (Qiagen, 74004). RNA quality was assessed using TapeStation (Agilent). The total RNA volume recovered was used for RNA-sequencing library preparation using SMART-Seq mRNA (Takara, 634773) and SMART-Seq Library Prep Kit (Takara, R400747) following the manufacturer’s instructions. cDNA amplification cycles were determined by quantitative PCR performed with 10% of the cDNA.

### RNA-sequencing and bioinformatic analysis

Paired-end sequencing of the libraries was performed on an AVITI^TM^ sequencer (Element Biosciences), with an average of 500 million reads per library. Samples were demultiplexed and initially analyzed using Galaxy^33^. Reads were trimmed to remove polyA sequences and adapters using Trimmomatic^34^. Trimmed reads were mapped to the zebrafish genome annotation GRCz12 using RNA STAR alignment tool^35^. Finally, gene-level read counts were obtained using featureCounts^36^. The resulting read count matrix was analyzed in R Studio (R version 4.5.2; Posit team (2025). RStudio: Integrated Development Environment for R. Posit Software, PBC, Boston, MA. URL: http://www.posit.co/). The BiocManager version used was 1.30.27. Differential gene expression analysis was performed using DESeq2^37^ (1.50.2). Genes were considered significantly differentially expressed using an adjusted p-value threshold of p_adjusted < 0.05, with a log_2_fold change threshold of > 0.5 for upregulated genes and < −0.5 for downregulated genes. Graphical representations were generated using the ggplot2 (4.0.1), enhancevolcano (1.28.2), and pheatmap (1.0.13) packages. Gene ontology enrichment analysis and gene set enrichment analysis (GSEA) were performed using ClusterProfiler^38^ (4.18.4) with annotation support from the org.Dr.eg.db (3.22.0) package. Significant enrichment was determined using an adjusted p-value cutoff of 0.05 with the FDR method.

### Statistics

All statistical analyses for experimental data were performed in GraphPad Prism 10. Distribution of data from all sample groups was assessed using the Shapiro-Wilks normality test. Comparative statistics between two sample groups was performed using the unpaired two-sided t-test for parametric data or the two-sided Mann–Whitney test for nonparametric data. Comparative statistics between more than two sample groups was performed using ordinary one-way ANOVA for parametric data or the Kruskal–Wallis test for nonparametric data. Specific multiple comparison tests used for experimental data with more than two sample groups can be found in the figure legends.

## RESULTS

### AP-1 regulates cardiomyocyte autophagy

To investigate the gene regulatory networks controlled by AP-1 transcription factors during the cardiomyocyte (CM) response to cardiac cryoinjury, we generated a new CM-specific dominant-negative A-Fos line in the safe harbor pIGLET locus^30^. Here, we removed EGFP fluorescence from the original line^8^, utilized a stronger ubiquitous promoter (ubbR)^27^ and inserted the transgene into the pIGLET14a locus to prevent epigenetic silencing of the transgene across generations (**Figure 1A**). Immunoblotting analysis of ventricles from *Tg(p14a.ubbR:loxP-Stop-loxP-FlagAFos-P2A-TagBFPHA); Tg(myl7:Cre-ERT2)* zebrafish (hereafter referred to as CM:AFos) revealed Flag-AFos protein in ventricles treated with tamoxifen and ethanol vehicle (**Supplementary Figure 1A**), indicating leakiness of the transgene in the absence of tamoxifen. Immunoblotting analysis of ventricles from *Tg(p14a.ubbR:loxP-Stop-loxP-FlagAFos-P2A-TagBFPHA)* zebrafish (hereafter referred to as control) revealed a lack of Flag-AFos protein in the absence of Cre recombinase (**Supplementary Figure 1A**). Immunostaining analysis confirmed CM specificity of CM:AFos (**Supplementary Figure 1B**). RNA-sequencing analysis of isolated CMs from control and CM:AFos uninjured ventricles and at 7 days post cryoinjury (dpci) revealed hundreds of differentially expressed genes (DEGs) when AP-1 transcription factor function is blocked (**Figure 1B and Supplementary Figures 1C and 1D**). Genes upregulated in CM:AFos CMs include interferon-inducible genes (*ifi27.6, ifi27.4*) and interferon regulatory factors (*irf1b, irf4b, irf4l*), immune activation, and apoptosis (*nlrp3, casp10, tnk1, bmf1*), indicating a hyperinflammatory state in CMs with blocked AP-1 transcription factor function (**Figure 1B**). Gene ontology (GO) and Kyoto Encyclopedia of Genes and Genomes (KEGG) pathway enrichment analysis revealed an enrichment in gene sets related to the innate immune response, chemokine activity, and cytokine receptor binding in CM:AFos isolated CMs compared to control from uninjured ventricles and at 7 dpci (**Supplementary Figures 1E and 1F**). Genes that were not upregulated in response to injury when CM AP-1 transcription factor function was blocked include genes associated with CM dedifferentiation and actin cytoskeleton organization (*mustn1b, tagln, cfl2, cnn1b, xirp2a*). These gene expression changes are in line with phenotypes we have previously observed in injured CM:AFos ventricles, including decreased CM dedifferentiation and protrusion of CMs into the injured area^8,31^. Additionally, we found several genes associated with the stress response that failed to upregulate in CM:AFos upon injury, including those involved in protein quality control and autophagy (*dnajb5, hspb9/hspb9l, fbxo40, sh3gl3b, pak1, fbxo32, ywhag2*) and redox homeostasis (*gsto1, mgst3a*) (**Figure 1B**). Furthermore, GO and KEGG pathway enrichment revealed a failure in the upregulation of gene sets involved in translation, cellular respiration, and oxidative phosphorylation when AP-1 transcription factor function is blocked in CMs (**Supplementary Figure 1E and 1F**). These data indicate that AP-1 promotes the expression of genes to allow CMs to cope with cellular stress in response to cardiac cryoinjury.

**Figure 1.**
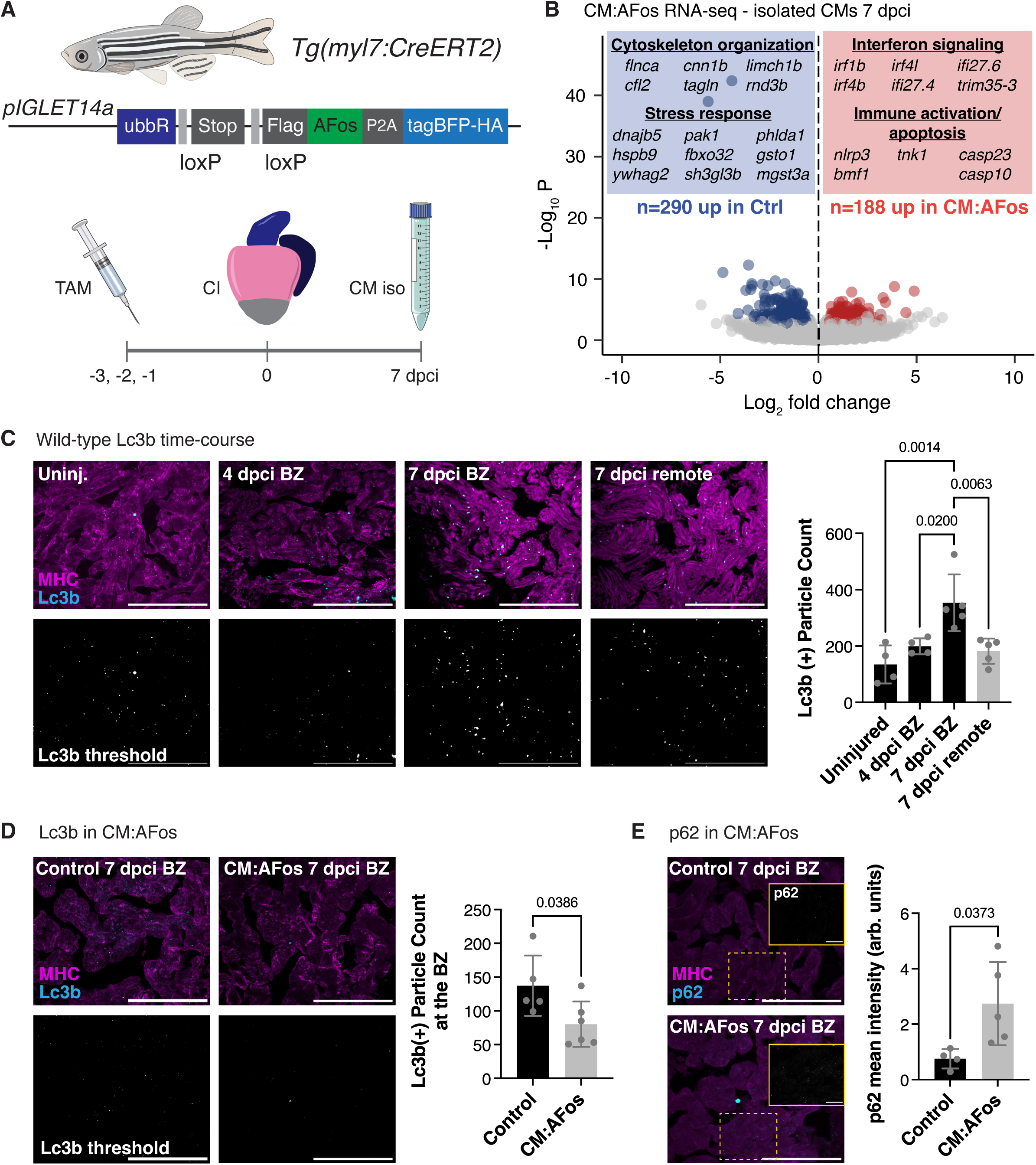
AP-1 transcription factors regulate cardiomyocyte autophagy during zebrafish heart regeneration. **(A)** Schematic illustrating *Tg(pIGLET14a.ubbR:loxP-Stop-loxP-FlagAFos-P2A-TagBFP-HA); Tg(myl7:Cre-ERT2)* line (hereafter referred to as CM:AFos) and the experimental scheme to isolate CMs expressing *AFos*. **(B)** Volcano plot depicting differentially expressed genes (DEGs) from bulk RNA-seq analysis of isolated CMs from control and CM:AFos ventricles at 7 dpci. Red and blue dots depict genes that are enriched in CM:AFos or control CMs, respectively (absolute(log_2_FC)>0.5, p_adj_<0.05). Red and blue boxes illustrate representative biological processes and genes enriched in CM:AFos and control CMs, respectively. **(C)** Immunostaining of MHC and Lc3b in cryosections of wild-type zebrafish uninjured (n=4) and cryoinjured ventricles at 4 dpci (n=4) and 7 dpci (n=5). Quantification of the number of Lc3b (+) autophagosome punctae at the border zone (BZ) is shown on the right. Thresholded Lc3b signal used for quantification is shown in the bottom panels. Data are presented as mean ± SD. P-value was calculated using one-way ANOVA with Tukey’s multiple comparisons test. **(D)** Immunostaining of MHC and Lc3b in cryosections of control (n=5) and CM:AFos (n=6) ventricles at 7 dpci. Quantification of the number of Lc3b (+) autophagosome punctae at the border zone (BZ) is shown on the right. Thresholded Lc3b signal used for quantification is shown in the bottom panels. Data are presented as mean ± SD. P-value was calculated using an unpaired two-tailed t-test. **(E)** Immunostaining of MHC and p62 (Sqstm1) in cryosections of control (n=4) and CM:AFos (n=5) ventricles at 7 dpci. Insets show zoomed in single channel images of p62 expression. Quantification of mean intensity of p62 is shown on the right. Data are presented as mean ± SD. P-value was calculated using an unpaired two-tailed t-test. Scale bar: 100 μm and 20 μm in insets in **(E)**.

We chose to focus our remaining analysis on AP-1 regulation of CM autophagy, based on a lack of clear consensus regarding the function of autophagy in zebrafish cardiac regeneration^23,24^ and the unknown function of CM-specific autophagy in this process. First, we determined that CMs at the border zone (BZ) exhibit Lc3b(+) punctae, likely corresponding to autophagosomes, at 7 dpci (**Figure 1C**) in line with previously published studies^23,24^. We observed increased enrichment of Lc3b(+) punctae at the wound BZ of regenerating zebrafish ventricles when compared to the remote zone (**Figure 1C**). Notably, we found fewer Lc3b(+) punctae at the wound BZ in CM:AFos ventricles compared to control at 7 dpci (**Figure 1D**). This decrease in Lc3b was accompanied by a corresponding increase in p62, indicative of a block in autophagy in CM:AFos ventricles compared to control (**Figure 1E**). Altogether, these results suggest that AP-1 promotes CM autophagy in response to injury in the adult zebrafish heart.

To test whether AP-1 promotes autophagy in mammalian CMs, we utilized neonatal rat ventricular myocyte (NRVM) culture. First, we treated NRVMs with cobalt chloride (CoCl_2_) to stabilize HIF-1α and pharmacologically mimic the hypoxia experienced by CMs in response to ischemic injury (**Figure 2A and 2B**). Notably, stabilization of HIF-1α and pharmacological mimicry of hypoxia led to an increase in levels of AP-1 family members JUN and FOSL2 (**Figure 2B**). We then tested whether stabilization of HIF-1α was sufficient to promote CM autophagy and found that levels of LC3-II increase further in response to CQ+CoCl_2_ treatment compared to CQ treatment alone, indicative of increased autophagic flux (**Supplementary Figure 2A**). Notably, pharmacological inhibition of AP-1 function through treatment of NRVMs with T-5224 blunted this increase in CM autophagy (**Figure 2C**). These data suggest that there may be an evolutionarily conserved role for AP-1 in promoting CM autophagy in response to injury and ischemia.

**Figure 2.**
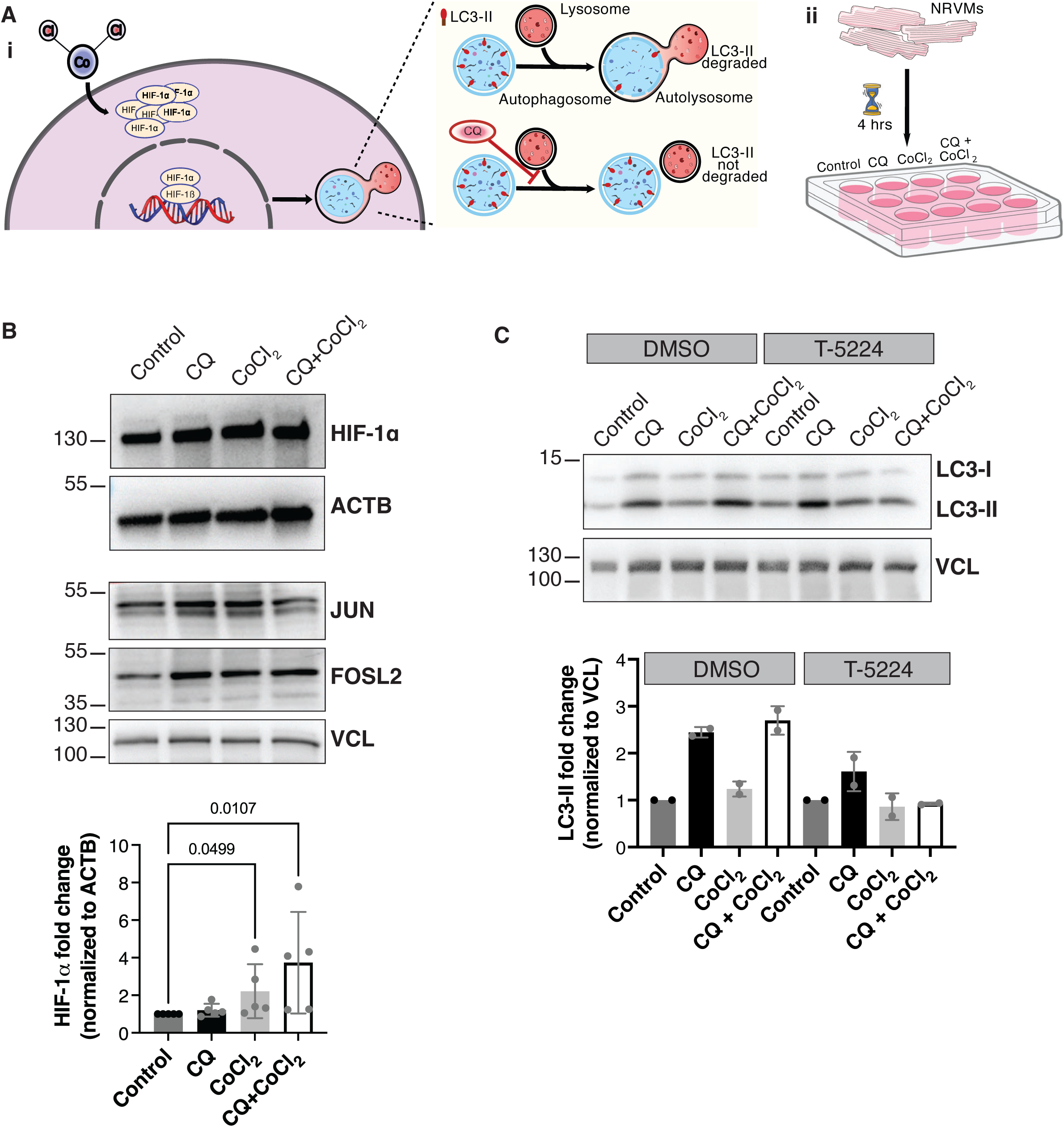
AP-1-mediated regulation of cardiomyocyte autophagy is conserved in mammalian cardiomyocytes. **(A)** Schematic illustrating effects of cobalt chloride (CoCl_2_) and chloroquine (CQ) on HIF-1α stabilization and autophagic flux, respectively (i). Treatment of neonatal rat ventricular myocytes (NRVMs) with CoCl_2_ and CQ to mimic hypoxia and promote and measure CM autophagy (ii). **(B)** Representative immunoblot of HIF-1α and AP-1 family members JUN and FOSL2 in control, CQ-treated, CoCl_2_-treated, and CQ+CoCl_2_-treated NRVMs. Quantification of HIF-1α/ACTB levels normalized to control NRVMs is shown in the graph on the bottom. Data are presented as mean ± SD. P-value was calculated using a Kruskal-Wallis test. **(C)** Representative immunoblot of LC3 and Vinculin (VCL) in control, CQ-treated, CoCl_2_-treated, and CQ+CoCl_2_-treated NRVMs in the presence of AP-1 inhibitor T-5224 or DMSO control. Quantification of LC3-II/VCL normalized to the respective controls (DMSO/T-5224) shown in the graph below. Data are presented as mean ± SD.

### Blocking cardiomyocyte autophagy impairs cardiac regeneration

Considering the pleiotropic effects of AP-1 transcription factor function in response to cardiac injury, and the unknown effects of CM autophagy on cardiac regeneration, we generated a genetic model to block autophagy in a CM-specific manner (**Figure 3A**). To this end, we generated a *Tg(p14a.ubbR:loxP-Stop-loxP-3xHAAtg4bC74A); Tg(myl7:Cre-ERT2)* zebrafish line (hereafter referred to as CM:Atg4bC74A) overexpressing mutant Atg4bC74A specifically in CMs. Mutant Atg4bC74A was previously shown to impair lipidation of LC3 and incorporation into autophagosomal membranes^17^. Further, expression of mutant *atg4bC74A* in zebrafish has been shown to inhibit autophagic flux *in vivo*^39^. As with the CM:AFos transgene, we observed leakiness of the CM:Atg4bC74A transgene in the absence of tamoxifen (**Figure 3B and Supplementary Figure 3A**) and utilized *Tg(p14a.ubbR:loxP-Stop-loxP-3xHAAtg4bC74A)* Cre (-) zebrafish as controls (hereafter referred to as Control). Blocking CM autophagy using the Atg4bC74A mutant resulted in a decrease in Lc3b(+) punctae at the border zone of CM:Atg4bC74A ventricles compared to control at 7 dpci (**Figure 3C**). This decrease in Lc3b was accompanied by a corresponding increase in p62 in CM:Atg4bC74A ventricles at 7 dpci (**Figure 3D**), suggesting that overexpression of Atg4bC74A blocks CM autophagy in zebrafish ventricles. To test whether blocking CM autophagy impairs cardiac regeneration, we performed Picrosirius red staining in ventricles from CM:Atg4bC74A and control zebrafish at 30 dpci. We observed an increase in the amount of collagen-containing scar tissue in CM:Atg4bC74A ventricles compared to control (**Figure 3E**), indicating the CM autophagy is important for resolution of the scar during cardiac regeneration in zebrafish.

**Figure 3.**
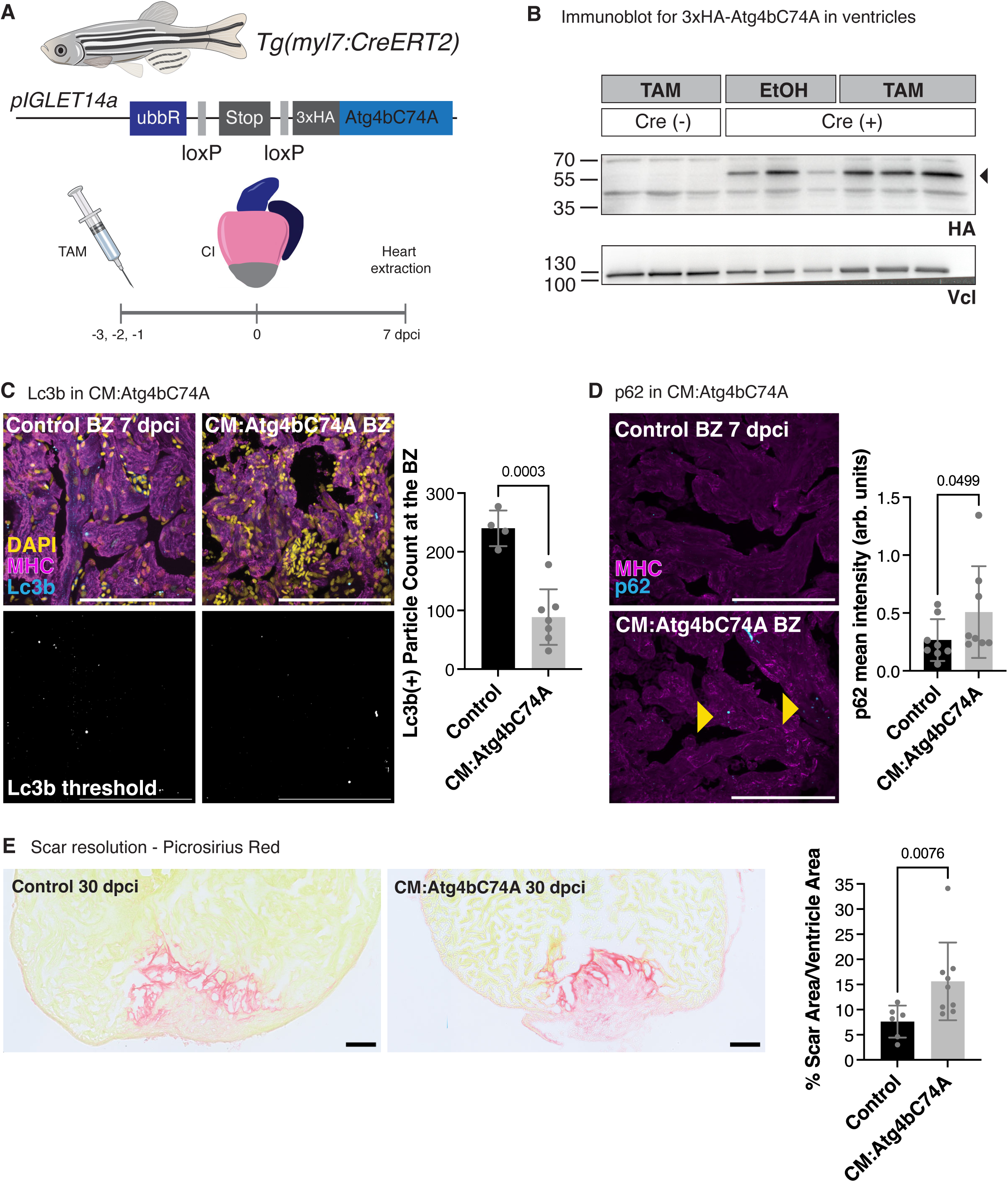
Cardiomyocyte-specific Atg4bC74A overexpression blocks autophagy and scar resolution in injured zebrafish hearts. **(A)** Schematic illustrating *Tg(pIGLET14a.ubbR:loxP-Stop-loxP-3xHAAtg4bC74A); Tg(myl7:Cre-ERT2)* line (hereafter referred to as CM:Atg4bC74A) and experimental scheme to induce CM-specific overexpression of *atg4bC74A*. **(B)** Immunoblot of ventricles from uninjured *Tg(pIGLET14a.ubbR:loxP-Stop-loxP-3xHAAtg4bC74A)* fish treated with tamoxifen (TAM), and CM:Atg4bC74A fish treated with ethanol (EtOH vehicle) or tamoxifen using antibodies for Vinculin (Vcl) and HA (n=3 biological replicates). Black arrowhead points to 3xHA-Atg4bC74A protein. **(C)** Immunostaining of DAPI, MHC and Lc3b in cryosections of ventricles from control (n=4) and CM:Atg4bC74A (n=7) zebrafish at 7 dpci. Quantification of the number of Lc3b (+) autophagosome punctae at the border zone (BZ) is shown on the right. Thresholded Lc3b signal used for quantification is shown in the bottom panels. Data are presented as mean ± SD. P-value was calculated using an unpaired two-tailed t-test. **(D)** Immunostaining of MHC and p62 in cryosections of ventricles from control (n=8) and CM:Atg4bC74A (n=8) zebrafish at 7 dpci. Yellow arrowheads denote p62 accumulation in CMs. Quantification of mean intensity of p62 is shown on the right. Data are presented as mean ± SD. P-value was calculated using a Mann-Whitney test. **(E)** Picrosirius red staining of collagen in ventricles from control (n = 6) and CM:Atg4bC74A (n =9) zebrafish at 30 dpci. Quantification of scar area (as a % of ventricle area) is shown on the right. Data are presented as mean ± SD. P-value was calculated using a Mann-Whitney test. Scale bar: 100 μm.

### Blocking cardiomyocyte autophagy affects cardiomyocyte invasion of injured tissue

To investigate how blocking CM autophagy leads to impairment in resolution of collagen-containing injured tissue, we tested several CM processes that are important for cardiac regeneration. First, we examined apoptosis at 24 hours post cryoinjury (hpci) using terminal deoxynucleotidyl transferase dUTP nick end labeling (TUNEL) and found no difference in the percentage of TUNEL+ nuclei in control and CM:Atg4bC74A ventricles at 24 hours post cryoinjury (hpci) (**Supplementary Figure 4A**). We assessed CM dedifferentiation and CM proliferation using immunostaining for embCMHC (N2.261) and PCNA/Mef2, respectively, and found no significant difference in CM:Atg4bC74A ventricles compared to control at 7 dpci (**Supplementary Figure 4B and 4C**). We then examined CM protrusion into the injured tissue at the wound border zone by staining of F-actin in thick cryosections (**Figure 4A**). We found that there was no difference in the number of CM protrusions at the wound border in CM:Atg4bC74A ventricles compared to control at 7 dpci, but a significant decrease in the length of protrusions (**Figure 4B**). As we have recently shown that macrophages play an important role in regulating CM protrusion length^31^, we analyzed the presence of macrophages at the wound border zone using a transgenic fluorescent *Tg(mpeg1:EGFP)* reporter line. Quantification of macrophages directly at the wound border zone revealed no significant change in macrophage number in CM:Atg4bC74A ventricles compared to control at 7 dpci (**Figure 4C**). Notably, while absolute numbers of macrophages did not change, we observed a decrease in phagocytic and pro-reparative macrophage markers, including *mrc1b* and Cxcr4b, within the wound and at the wound border zone in CM:Atg4bC74A ventricles compared to control at 7 dpci (**Figure 4D and Supplementary Figure 4D**). The presence of neutrophils and localization of the pro-inflammatory cytokine Tnfa at 24 hours post cryoinjury (hpci) were not affected in CM:Atg4bC74A ventricles compared to control (**Supplementary Figure 4E and 4F**). Altogether, these results indicate that blocking CM autophagy affects macrophage phenotype, without affecting the initial recruitment of neutrophils, upon cardiac injury.

**Figure 4.**
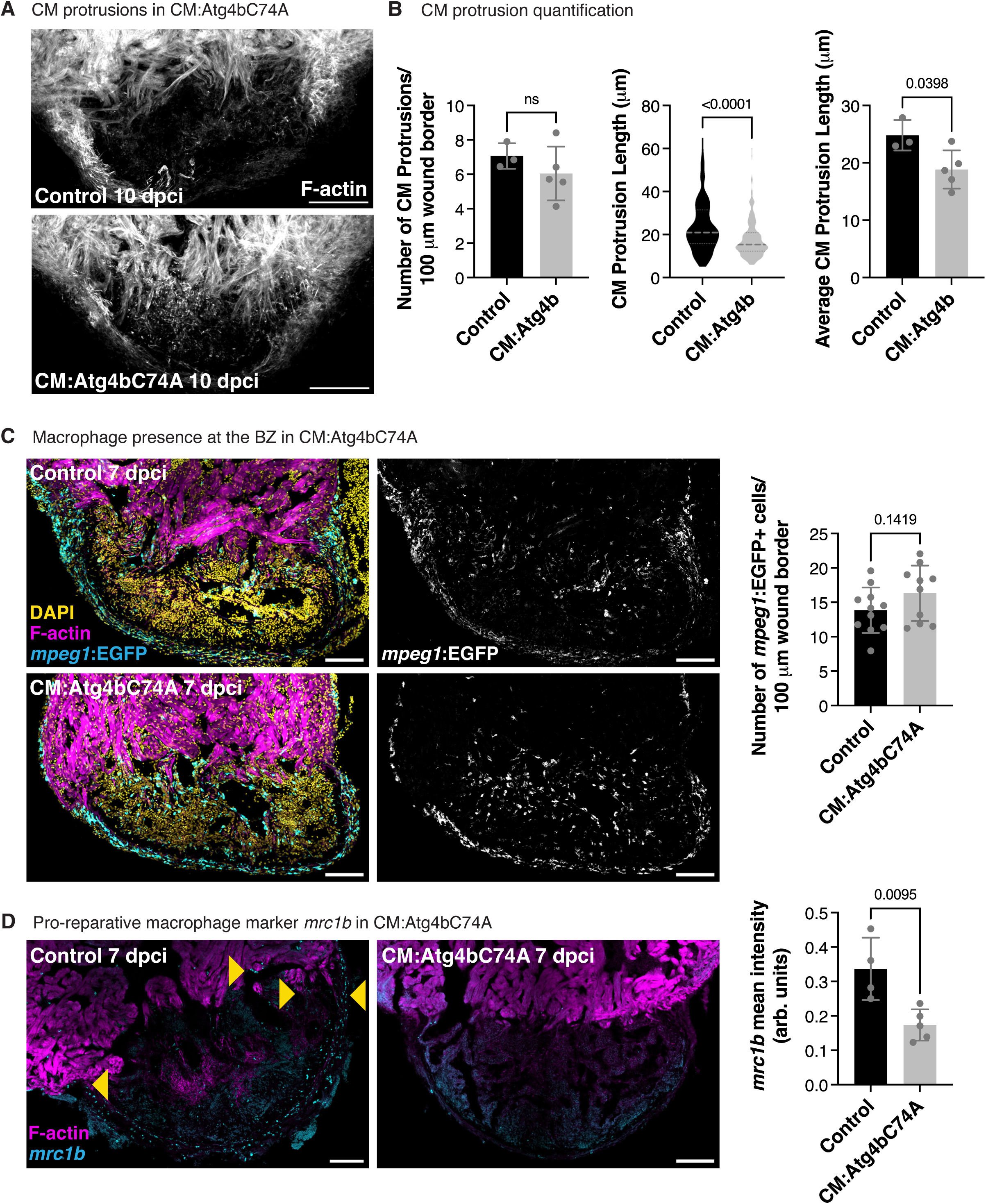
Blocking cardiomyocyte autophagy affects cardiomyocyte protrusion into the injured tissue. **(A)** F-actin staining of thick cryosections in ventricles from control (n=3) and CM:Atg4bC74A (n=5) zebrafish at 10 dpci. **(B)** Quantification of the number of CM protrusions per 100 μm wound border is shown on the left. Data are presented as mean ± SD. P-value was calculated using an unpaired two-tailed t-test. Quantification of CM protrusion length is shown in the middle. Data are presented as violin plots of all points with thick dotted gray line indicating the median and thin dotted gray lines indicating 25^th^ and 75^th^ percentile. P-values were calculated using a Mann-Whitney test. Quantification of the average CM protrusion length per ventricle is shown on the right. Data are presented as mean ± SD. P-value was calculated using an unpaired two-tailed t-test. **(C)** Immunostaining of GFP, F-actin and DAPI in ventricles from control (n=11) and CM:Atg4bC74A (n=10) *Tg(mpeg1:GFP)* zebrafish at 7 dpci. Quantification of the number of *mpeg1*:EGFP+ cells per 100 μm of wound border is shown on the right. Data are presented as mean ± SD. P-value was calculated using an unpaired two-tailed t-test. **(D)** F-actin staining and HCR *in situ* hybridization of *mrc1b* in ventricles from control (n=4) and CM:Atg4bC74A (n=5) zebrafish at 7 dpci. Yellow arrowheads denote *mrc1b*+ cells at the wound BZ. Quantification of mean intensity of *mrc1b* signal is shown on the right. Data are presented as mean ± SD. P-value was calculated using an unpaired two-tailed t-test. Scale bar: 100 μm

### Disrupting cardiomyocyte autophagy influences cardiac macrophage phenotype

To further investigate the phenotype of cardiac macrophages in the absence of CM autophagy, we performed bulk RNA-seq of macrophages isolated by Csf1r antibody-mediated fluorescence activated cell sorting (**Figure 5A and Supplementary Figure 5A and 5B**). Comparison of the transcriptome of macrophages from CM:Atg4bC74A and control ventricles at 7 dpci revealed 367 DEGs (259 upregulated, 108 downregulated, **Figure 5B**). Gene ontology and KEGG pathway enrichment analysis revealed a downregulation of genes associated with the lysosome and proteasome (**Supplementary Figure 5C**), in line with the decrease in *mrc1b* HCR signal observed in CM:Atg4bC74A ventricles compared to control at 7 dpci. Gene ontology and KEGG pathway enrichment analysis revealed an upregulation of genes associated angiogenesis, extracellular matrix (ECM) organization, muscle cytoskeleton, and focal adhesion/integrin signaling pathway in macrophages isolated from CM:Atg4bC74A ventricles compared to control (**Supplementary Figure 5C**). Examination of the expression of genes encoding canonical zebrafish macrophage markers (*csf1ra, cd68, mfap4.1, mpeg1.1, ccr2, itgam.1*) and cytokines associated with pro- (*tnfa, il6, il1b*) and anti-inflammatory (*il10, cxcr4b*) macrophages revealed few significant changes in the expression of pro-/anti-inflammatory markers (**Figure 5C**). Consistent with our HCR in situ hybridization data, phagocytic macrophage marker genes (*mrc1b*) were significantly downregulated, while pro-fibrotic marker genes (*tgfb2*) were upregulated in macrophages isolated from CM:Atg4bC74A ventricles compared to control at 7 dpci (**Figure 5C**). Gene set enrichment analysis confirmed that there was a significant decrease in gene expression signatures related to the inflammatory response, while gene expression signatures related to extracellular matrix (ECM) and angiogenesis were significantly upregulated (**Figure 5D**). Altogether, these analyses indicate that CM autophagy is important for the phenotypic state of cardiac macrophages in response to injury.

**Figure 5.**
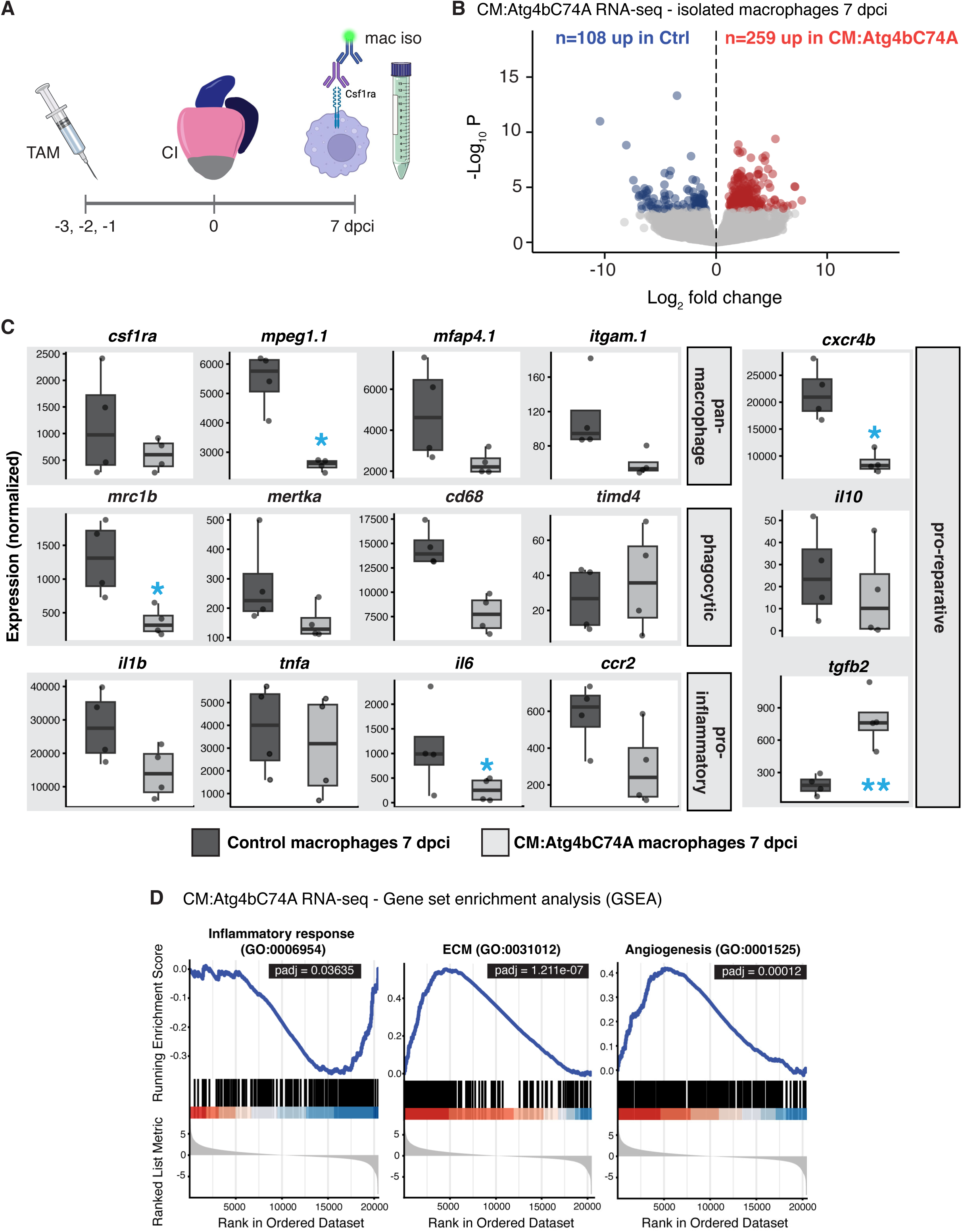
Blocking cardiomyocyte autophagy affects the transcriptome of cardiac macrophages following injury. **(A)** Experimental scheme to induce CM-specific *atg4bC74A* expression and isolate macrophages from cryoinjured ventricles for RNA-sequencing. **(B)** Volcano plot depicting differentially expressed genes (DEGs) from bulk RNA-seq analysis of isolated macrophages from control and CM:Atg4bC74A ventricles at 7 dpci. Red and blue dots depict genes that are enriched in CM:Atg4bC74A or control CMs, respectively (absolute(log_2_FC)>0.5, p_adj_<0.05). **(C)** Box and whisker plots of selected genes representing pan-macrophage, phagocytic, pro-inflammatory, and pro-reparative macrophage marker genes. Blue asterisks denote genes with absolute(log_2_FC)>0.5 and p_adj_<0.05. **(D)** Gene set enrichment analysis of bulk RNA-seq analysis of macrophages isolated from control and CM:Atg4bC74A ventricles at 7 dpci. The blue line denotes the running enrichment score, and the black lines in the x-axis represent the distribution of the gene markers across the ranked gene list based on log2FC (bottom plot). The p-value was corrected with the FDR method.

Next, we visualized the expression of collagen genes in macrophages based on the enrichment of ECM gene ontology in CM:Atg4bC74A ventricles and previously published data showing that a subset of macrophages deposit collagen into the wound during zebrafish cardiac regeneration^40^. Heat map representation of RNA-seq revealed increased collagen gene expression in macrophages isolated from CM:Atg4bC74A ventricles compared to control at 7 dpci (**Figure 6A**). Staining for fibrillar collagen revealed an increase in collagen protein levels in CM:Atg4bC74A ventricles compared to control at 7 dpci, in line with the bulk RNA-seq analysis from isolated macrophages (**Figure 6B**). Further confirmation using HCR *in situ* hybridization for *col1a1a* and co-localization analysis with *mpeg1:*EGFP+ cells in ventricular tissue sections revealed an increase in the ratio of *col1a1a*+ macrophages and an increase in *col1a1a* mean intensity within GFP+ macrophages in CM:Atg4bC74A ventricles compared to control (**Figure 6C**). Together, these results indicate that blocking CM autophagy shifts macrophage phenotype from a pro-inflammatory/phagocytic to a pro-fibrotic state.

**Figure 6.**
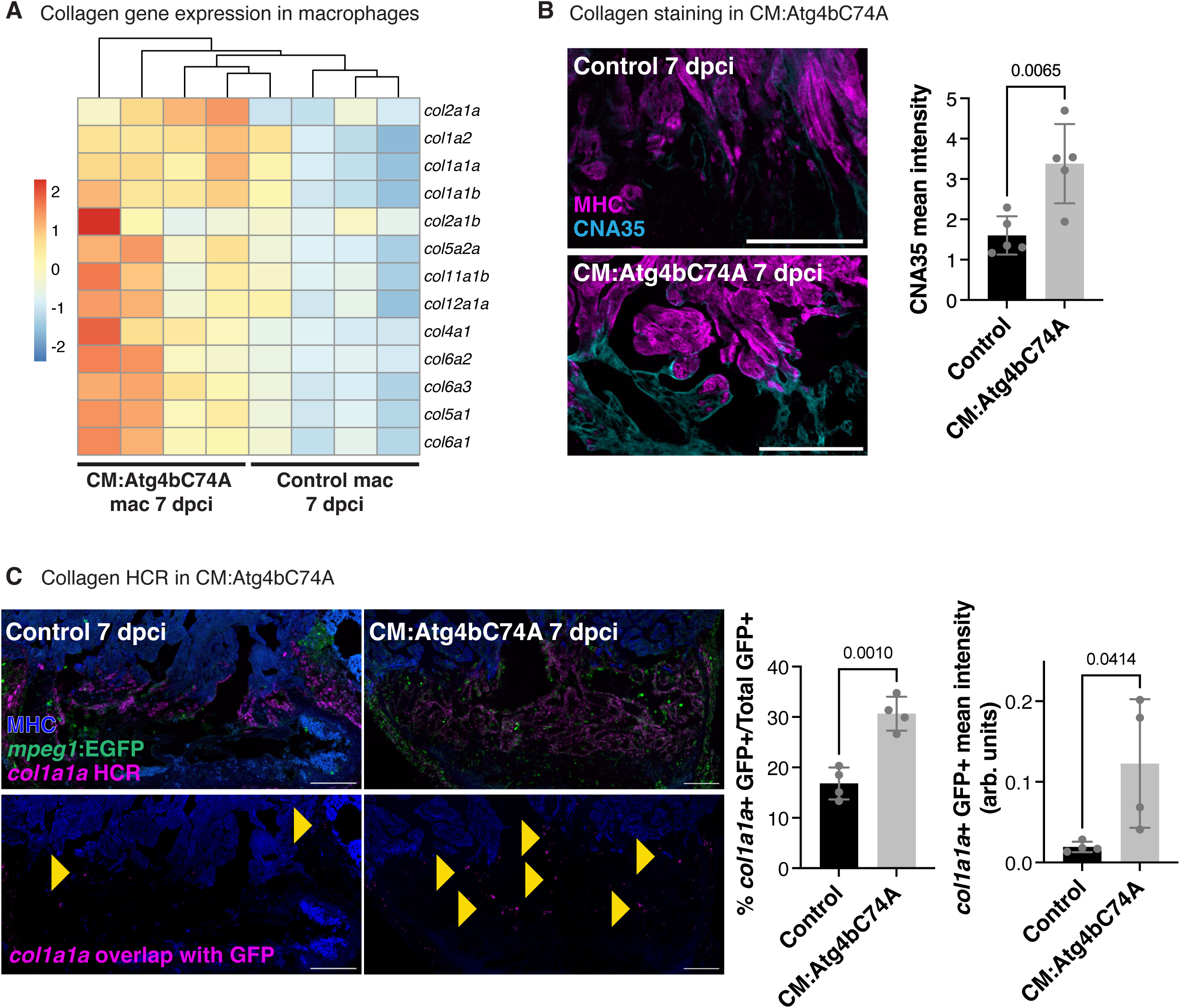
Collagen gene expression is upregulated in macrophages when CM autophagy is blocked. **(A)** Heat map of collagen gene normalized expression (row) from bulk RNA-seq in macrophages isolated from control and CM:Atg4bC74A ventricles at 7 dpci (column). **(B)** F-actin and fibrillar collagen (CNA35) staining in ventricles from control (n=5) and CM:Atg4bC74A (n=5) zebrafish at 7 dpci. Quantification of CNA35 mean intensity is shown on the right. Data are presented as mean ± SD. P-value was calculated using an unpaired two-tailed t-test. **(C)** MHC/GFP immunostaining and HCR *in situ* hybridization of *col1a1a* in ventricles from control (n=4) and CM:Atg4bC74A (n=5) *Tg(mpeg1:EGFP*) zebrafish at 7 dpci. Yellow arrowheads point to *col1a1a* signal that overlaps with *mpeg1*:EGFP+ cells. Quantification of *col1a1a*/*mpeg1*:EGFP+ cells as a ratio of total *mpeg1*:EGFP+ cells, and mean intensity of *col1a1a* signal colocalized with *mpeg1*:EGFP+ is shown on the right. Data are presented as mean ± SD. P-value was calculated using an unpaired two-tailed t-test. Scale bar: 100 μm

## DISCUSSION

Using newly generated genetic tools, we show here that CM autophagy is important for CM protrusion into and resolution of the injured tissue during zebrafish cardiac regeneration. Our results are consistent with previous reports showing a cardioprotective role for CM autophagy in response to permanent coronary artery occlusion and myocardial ischemia^19^. However, the role of CM autophagy is complex and context-dependent, and autophagy has been shown to promote maladaptive responses to disease and injury models, including ischemia/reperfusion and pressure overload^20–22,41^. Notably, in zebrafish cardiac regeneration, there have been two studies on the role of autophagy that differ in their conclusions as to whether autophagy promotes or inhibits regeneration^23,24^. Our study provides evidence for a pro-regenerative role of CM autophagy upon cardiac cryoinjury.

Strikingly, we show that blocking CM autophagy had no effect on cell survival, CM dedifferentiation or proliferation, but rather on the transcriptomes of cardiac macrophages within the injured heart. Our results align with a previous landmark study showing that cardiac macrophages help maintain homeostasis in the heart through phagocytosis of defective mitochondria extruded from CMs, so-called CM exophers^42^. Notably, the authors show that macrophage uptake of defective mitochondria derived from CMs increased in conditions of cardiac stress/injury and was dependent on CM autophagy. The results of our studies here add further insight into the phenomenon of CM:macrophage crosstalk in response to cardiac injury: CM autophagy regulates the presence of phagocytic macrophages and limits pro-fibrotic gene expression signatures upon cardiac injury. This crosstalk likely occurs through direct exchange of cellular material, as we and others have observed macrophage uptake of fluorescent CM membrane material in zebrafish hearts^31,43^. Recent studies in the neonatal murine heart and the zebrafish spinal cord indicate that macrophage-mediated phagocytosis/efferocytosis is an important regulator of regeneration^44,45^. How CM autophagy affects phagocytic uptake of CM cellular material and the mechanisms underlying its effect on macrophage transcriptomic signatures, however, remain to be investigated.

We show here that AP-1 promotes CM autophagy in response to injury and stress. We and others have shown that AP-1 is an essential regulator of cardiac regeneration in zebrafish through its ability to promote changes in chromatin accessibility and at putative enhancer regions^8,9^. Notably, AP-1 TF binding sites motifs have also been found in chromatin that becomes more accessible in non-regenerative CMs^46^. Functional analysis of regeneration-responsive enhancers has suggested that AP-1 motifs are an ancestral component of enhancers that promote gene expression programs to activate regeneration, which includes both an injury response and regenerative output^47^. The authors suggest that in regeneration-incompetent animals, these ancestral enhancers may have been repurposed and led to the retention of injury response without regenerative output. We speculate that the regulation of CM autophagy by AP-1 may represent a conserved component of the CM injury response in teleosts and mammals. However, AP-1 controls additional aspects of the CM regeneration program, and while CM autophagy constitutes an important part of the regenerative response, it is insufficient to confer regenerative capacity in nonregenerative CMs.

## ACKNOWLEDGEMENTS

The authors thank the Mosimann and Stainier labs for sharing zebrafish lines and the pT3TS-phiC31 mRNA plasmid. The authors gratefully acknowledge the data storage service SDS@hd supported by the Ministry of Science, Research, and the Arts Baden-Württemberg (MWK) and the German Research Foundation (DFG) through grant INST 35/1803-1 FUGG and INST 35/1804-1 LAGG. The authors acknowledge the support of the Freiburg Galaxy Team: Bioinformatics, University of Freiburg (Germany), funded by the German Federal Ministry of Education and Research BMFTR grant 031 A538A de.NBI-RBC and the Ministry of Science, Research and the Arts Baden-Württemberg (MWK) within the framework of LIBIS/de.NBI Freiburg.

