## Supplementary Figures for "Cardiomyocyte autophagy promotes a pro-regenerative immune response during cardiac regeneration"

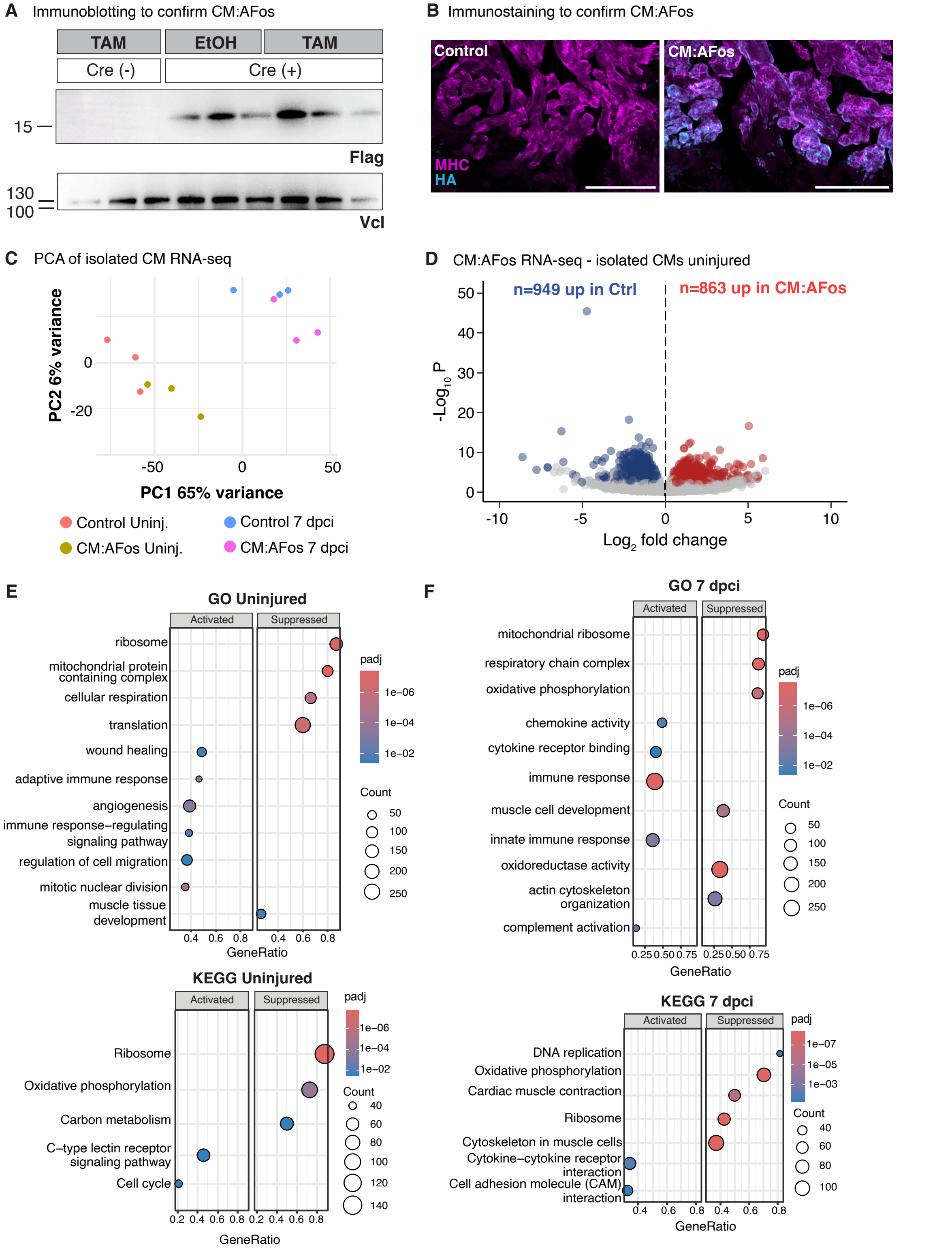


**Supplementary Figure 1. CM:AFos RNA-seq reveals targets of AP-1 in response to cardiac injury. (A)** Immunoblotting with anti-Flag and Vinculin (Vcl) antibodies to test the presence of Flag-AFos protein in ventricles from *Tg(p14a.ubbR:loxP-Stop-loxP-FlagAFos-P2A-tagBFP-HA)* Cre(-) (n=3) and *Tg(p14a.ubbR:loxP-Stop-loxP-FlagAFos-P2A-tagBFP-HA); Tg(myl7:Cre-ERT2)* zebrafish treated with tamoxifen (TAM, n=3) and vehicle (EtOH, n=3). **(B)** MHC and HA immunostaining to confirm localization of Flag-AFos-P2A-tagBFP-HA in cardiomyocytes in control and CM:AFos ventricles. **(C)** Principal component analysis of bulk RNA-seq of isolated CMs from control and CM:AFos uninjured and 7 dpci ventricles. Each dot represents an individual sample. **(D)** Volcano plot depicting differentially expressed genes (DEGs) from bulk RNA-seq analysis of isolated CMs from control and CM:AFos uninjured ventricles. Red and blue dots depict genes that are enriched in CM:AFos or control CMs, respectively (absolute($log_2FC$)>0.5, $p_{adj}<0.05$). **(E,F)** Dot-plot of selected significant ($p_{adj}<0.05$) gene ontology and KEGG pathway enrichment in bulk RNA-seq analysis of isolated CMs from control and CM:AFos uninjured (E) and 7 dpci (F) ventricles. The p-value was corrected using the FDR method. The size of the dot represents the number of genes and the color of the dot represents the value of the adjusted p-value. Scale bar: 100 μm

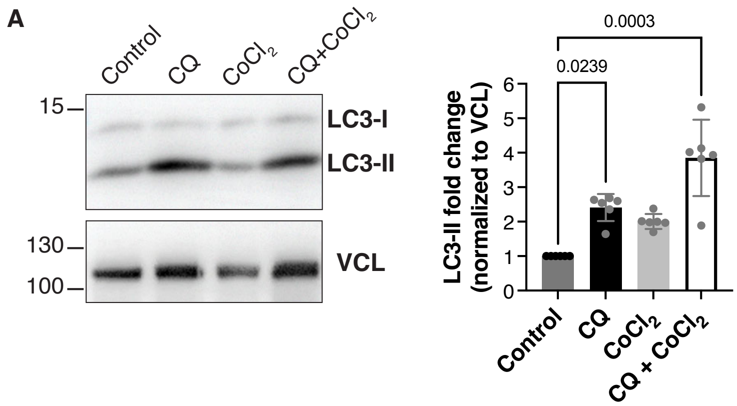


**Supplementary Figure 2. Stimulation of autophagy in neonatal rat ventricular myocytes.** **(A)** Representative immunoblot of LC3 (LC3-I and LC3-II) and Vinculin (VCL) in neonatal rat ventricular myocytes (NRVMs) treated with chloroquine (CQ), cobalt chloride ($CoCl_2$), and CQ+ $CoCl_2$. Quantification of LC3-II/VCL normalized to control NRVMs shown in the graph below. Data are presented as mean ± SD. P-value was calculated using an unpaired two-tailed t-test.

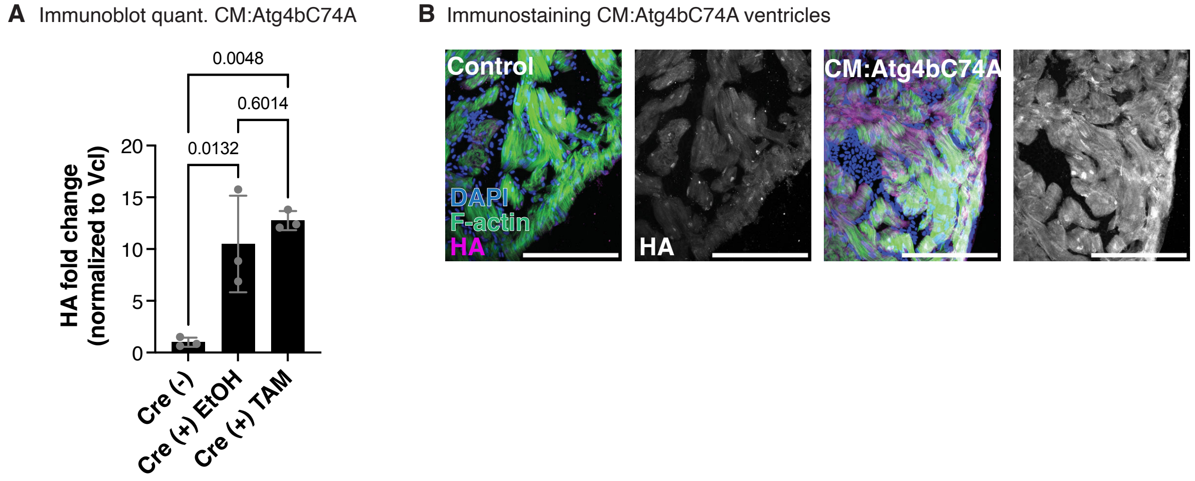


**Supplementary Figure 3. Characterization of CM:Atg4bC74A ventricles. (A)** Quantification of HA fold change (normalized to Vcl) from the immunoblot in Figure 3B. Data are presented as mean ± SD. P-value was calculated using one-way ANOVA with Tukey's multiple comparison test. **(B)** F-actin and DAPI staining, and HA immunostaining in uninjured control and CM:Atg4bC74A ventricles. Scale bar: 100 μm

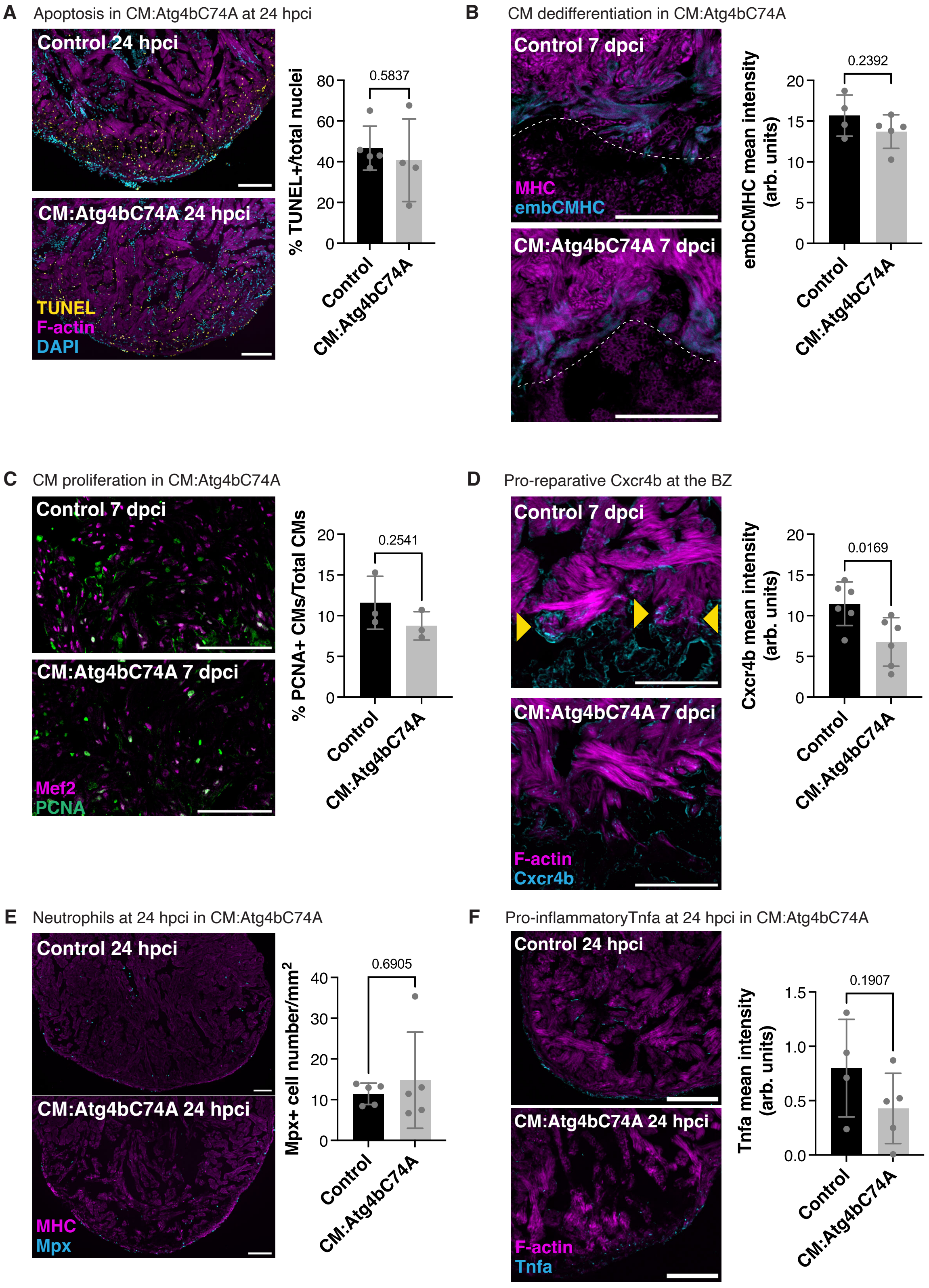


**Supplementary Figure 4. Characterization of CM regeneration parameters in CM:Atg4bC74A ventricles. (A)** TUNEL, F-actin, and DAPI staining in control (n=5) and

CM:Atg4bC74A (n=4) ventricles at 24 hours post cryoinjury (hpci). Quantification of % TUNEL+/total nuclei is shown on the right. Data are presented as mean ± SD. P-value was calculated using an unpaired two-tailed t-test. **(B)** embCMHC (N2.261) and MHC immunostaining in control (n=4) and CM:Atg4bC74A (n=5) ventricles at 7 dpci. Quantification of embCMHC mean intensity is shown on the right. Data are presented as mean ± SD. P-value was calculated using an unpaired two-tailed t-test. **(C)** Mef2 and PCNA immunostaining in control (n=3) and CM:Atg4bC74A (n=3) ventricles at 7 dpci. Quantification of PCNA+ CMs/total CMs within 100 μm of the wound border is shown on the right. Data are presented as mean ± SD. P-value was calculated using an unpaired two-tailed t-test. **(D)** Cxcr4b immunostaining and F-actin staining in control (n=6) and CM:Atg4bC74A (n=6) ventricles at 7 dpci. Yellow arrowheads denote Cxcr4b+ cells interacting with BZ CMs. Quantification of Cxcr4b mean intensity at the border zone is shown on the right. Data are presented as mean ± SD. P-value was calculated using an unpaired two-tailed t-test. **(E)** MHC and Mpx immunostaining in control (n=5) and CM:Atg4bC74A (n=5) ventricles at 24 hpci. Quantification of Mpx+ cell number is shown on the right. Data are presented as mean ± SD. P-value was calculated using a Mann-Whitney test. **(F)** F-actin staining and Tnfa immunostaining in control (n=4) and CM:Atg4bC74A (n=5) ventricles at 24 hpci. Quantification of Tnfa mean intensity is shown on the right. Data are presented as mean ± SD. P-value was calculated using an unpaired t-test. Scale bar: 100 μm

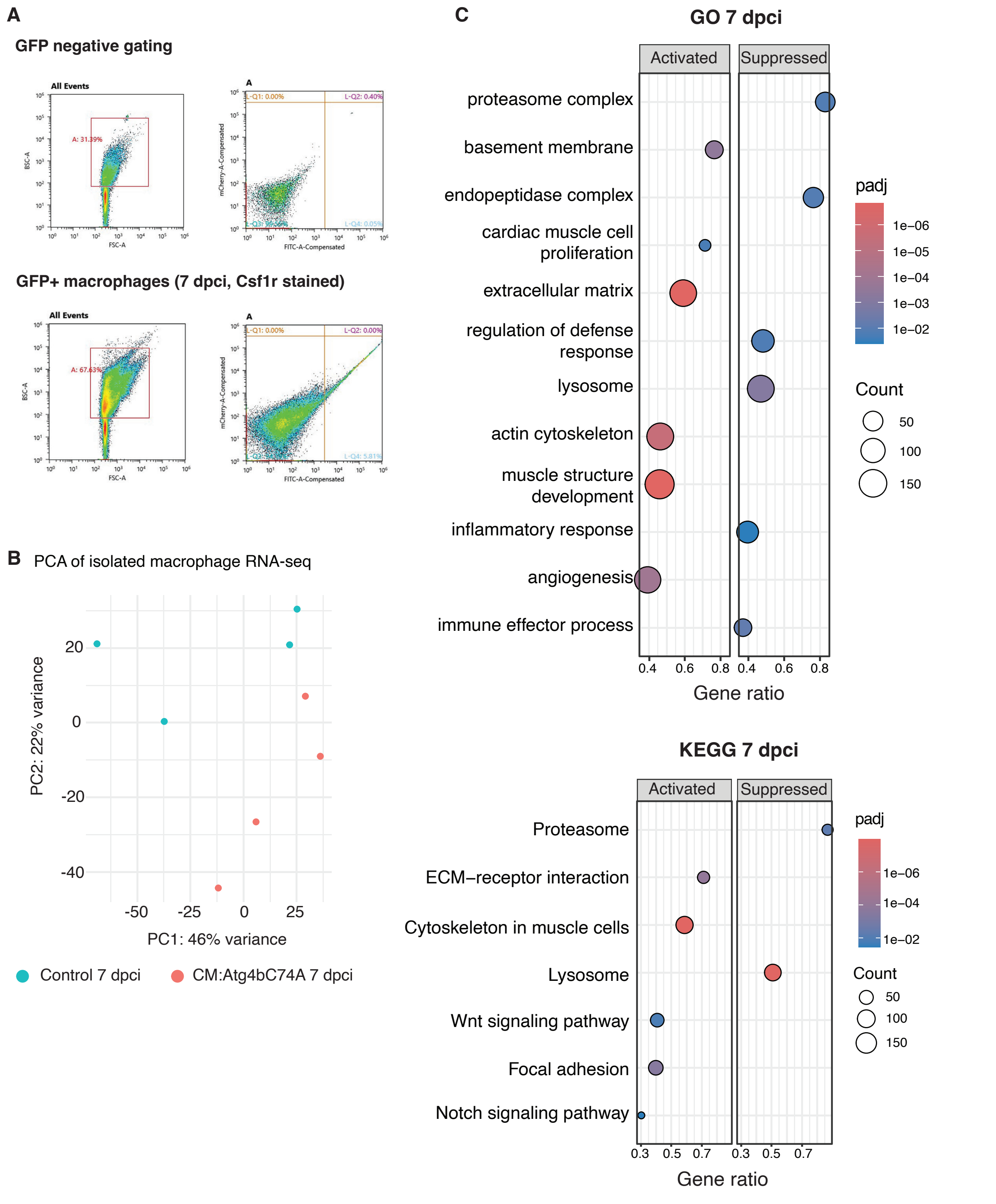


**Supplementary Figure 5. Bulk RNA-sequencing of macrophages isolated from CM:Atg4bC74A ventricles. (A)** Plots showing the hierarchical gating strategy for FACS-sorting live GFP+ macrophages and excluding non-fluorescent cells. **(B)** Principal component analysis of bulk RNA-seq of isolated macrophages from control and CM:Atg4bC74A ventricles at 7 dpci.

Each dot represents an individual sample. **(C)** Dot-plot of selected significant ($p_{adj}$<0.05) gene ontology and KEGG pathway enrichment in bulk RNA-seq analysis of isolated macrophages from control and CM:Atg4bC74A ventricles at 7 dpci. The p-value was corrected using the FDR method. The size of the dot represents the number of genes and the color of the dot represents the value of the adjusted p-value.
